# Evidence of subjective, but not objective, cognitive deficit in new mothers at one-year postpartum

**DOI:** 10.1101/2021.06.07.447303

**Authors:** Edwina R Orchard, Phillip GD Ward, Gary F Egan, Sharna D Jamadar

## Abstract

The experience and even existence of cognitive deficits in the postpartum period is uncertain, with only a few scientific studies, reporting inconsistent results. Here we investigate cognition in 86 women (43 first-time mothers one year postpartum, and 43 non-mothers). Mothers and non-mothers showed no significant differences on measures of objective cognition (verbal memory, working memory, processing speed or theory of mind). Despite the absence of objective differences, mothers self-reported significantly worse subjective memory than non-mothers. To interpret the difference between objective and subjective measures of memory, we investigated relationships between subjective memory, objective memory, and wellbeing. Mothers, but not non-mothers, showed a positive correlation between subjective and objective measures of memory, indicating mothers are ‘in-tune’ with their memory performance. Mothers also demonstrated a positive relationship between subjective memory and wellbeing (sleep, anxiety and depression), where better wellbeing correlated with higher subjective memory. This relationship was not apparent in non-mothers. The results suggest that poorer sleep, higher anxiety and higher depression are related to reports of poorer self-reported memory in mothers. Our results add to our growing understanding of maternal cognition at one year postpartum, with no evidence of cognitive differences between mothers and non-mothers.

---

‘Baby-brain’, also known as ‘pregnancy-brain’, or ‘mum-brain’, is a colloquial term that refers to cognitive decline experienced by women during the transition to motherhood (pregnancy and the postpartum period), and is self-reported by approximately 80% of expectant mothers^1^. Despite the overwhelming number of anecdotal reports of forgetfulness in pregnancy, empirical research has revealed that cognitive deficits in pregnancy are objectively subtle and remain within the normal ranges of general cognitive functioning and memory, noticeable to women themselves, but unlikely to cause significant disruption to daily life (e.g. reduced job performance)^1,2^. In addition to the question of whether cognitive deficits exists in the perinatal period, it is also unknown *when* memory deficits might cease – do pregnancy-related cognitive deficits end at child-birth? Or do they continue within the postpartum period?

Subtle memory deficits have been consistently found during pregnancy^1,3,4^, including objective deficits in working memory, recall, processing speed, and general cognition, as well as decrements in ratings of subjective memory. However, there are far fewer studies investigating cognition in the postpartum period, and evidence for differences in cognitive performance between postpartum versus non-postpartum women is mixed. When assessing the extant literature as a whole, it is clear that the number of studies showing non-significant changes or differences in cognitive function in the postpartum period^5–18^ outweigh those showing significant cognitive deficits^19–22^, relative to women outside the postpartum period.

The endurance of cognitive deficits into the postpartum period therefore remains unclear in the scientific literature. Poor understanding about the time-course of cognitive changes in early motherhood represents a shortcoming in maternal healthcare, and has implications for how women’s cognitive abilities in the postpartum period are perceived by society, employers, and themselves. One challenge to understanding the presence and magnitude of cognitive changes across the postpartum period is that early motherhood is a dynamic period in a woman’s life, involving considerable physiological and environmental changes. For example, mothers experience extreme fluctuations in sex steroid hormones, and significant alterations to sleep and mood across the peripartum period, which are known to impact cognitive ability^23–27^. These physiological and environmental experiences change dynamically across the postpartum period, being highly variable in the early postpartum, and stabilising in later postpartum months^28–30^. Thus, cognitive results obtained in the early first few weeks or months postpartum may not accurately predict cognitive outcome in the late postpartum period, or beyond.

One important question is how long pregnancy-related cognitive changes persist into the postpartum period. Women often report subjective experiences of cognitive difficulties and ‘brain-fog’ related to pregnancy and parenthood that do not necessarily resolve upon the birth of the child^7,8,12,14,31,32^, and postpartum mothers have shown objective deficits in verbal memory, processing speed, spatial recognition, and executive function^19–22^.

Thus, the goal of the current study was to investigate whether subjective and objective differences in cognition between mothers and non-mothers are evident late in the postpartum period. To this end, we measured differences in subjective and objective cognition between first-time mothers (one-year postpartum), and age- and education-matched women who had never been pregnant. Consistent with the anecdotal and self-report literature^7,8,12,14,31,32^, we hypothesised that mothers would self-report worse subjective memory than non-mothers but out-perform non-mothers in objective tests of cognitive performance (verbal memory, working memory, processing speed, and theory of mind). To foreshadow the results, mothers differed significantly from non-mothers on subjective, but not objective, measures of cognition. To explore these unexpected results, we undertook two *post-hoc* analyses; firstly to explore the alignment, or fidelity, between objective and subjective measures of cognition in mothers and non-mothers. Secondly, to explore the relationship between postpartum cognition and maternal wellbeing.

## Methods

### Participants

Our sample comprised 86 women, 43 first-time mothers approximately one-year postpartum (Mean±SD=11.6±1.7 months) and 43 women who had never been pregnant (for longer than eight weeks). Eleven mothers and one non-mother had a previous pregnancy that did not result in live birth (less than eight weeks gestation). All participants were aged over 18 years and right-handed. Inclusion criteria for the study included fluency in English, not currently pregnant (all participants completed a pregnancy test), no history of head injury, neurological disorder, epilepsy or stroke, and completion of an MRI safety survey for subsequent MRI scanning (e.g. presence of metal in the body, past surgeries, and claustrophobia). MRI results are not reported here.

### Procedure

After providing informed consent, all participants completed a psychosocial and cognitive battery, including paper and computer-based tasks and questionnaires. Demographic information and medical and obstetric history (including information about pregnancy, birth, breast-feeding, and time spent in primary care) was collected via clinical interview. Cognitive measures included the Hopkins Verbal Learning Test (HVLT; verbal memory)^33^, Digit Span Forward (short-term memory) and Backward^34^ (working memory), Symbol-Digit Modalities Task (SDMT; processing speed)^35^; Reading the Mind in Films Task (RMFT; Theory of Mind)^36^, and the Prospective and Retrospective Memory Questionnaire (PRMQ; subjective memory)^37^. The PRMQ measures self-reported memory errors across six subscales, reflecting different aspects of memory, including prospective and retrospective memory, short-term and long-term memory, and self- and environmentally-cued memory. Psychosocial measures included the Pittsburgh Sleep Quality Index (PSQI)^38^, Beck Depression Inventory (BDI)^39^, Beck Anxiety Inventory (BAI)^40^, Maternal Postnatal Attachment Scale (MPAS)^41^, and Maternal Self-Efficacy Questionnaire (MEQ)^42^. The tests and tasks we selected are commonly used in studies of motherhood to assess the cognitive domains tested here (Casey, 2000; Christensen et al., 2010; Glynn, 2010; Logan et al., 2014; Stark, 2000; Harris et al., 1996; Swain et al., 1997; Cuttler et al., 2011). Detailed descriptions of these tasks and questionnaires can be found in the Supplement. Participants completed the PRMQ, BDI, BAI, MPAS, and MEQ questionnaires at a time of their convenience on their own devices (e.g. laptop, phone) via a Qualtrics survey, before attending their testing session. This protocol was approved by the Human Research Ethics Committee (Project ID: 19455) and conducted in accordance with the relevant guidelines.

### Data Analysis

All 86 participants completed all tasks and questionnaires, with no missing data. Variables that were not normally distributed (HVLT recall, HVLT delayed recall, Digit Span Forward and Backward, SDMT, RMFT, PRMQ, PSQI, BDI, BAI) were log-transformed before further analysis. Histograms showing the frequency distributions of scores on the cognitive tasks are shown in Supplementary Figure 1. The HVLT delayed recall score showed evidence of ceiling effects, and was not included for further analysis, though results of the group comparison was included for completeness. Comparisons between groups were completed with independent samples *t*-tests, with alpha set to .05 as a threshold for statistical significance, and effect sizes were calculated with Cohen’s *d*^43^. Multiple comparisons were Bonferroni-corrected by dividing alpha by the number of tests run (e.g. for six tests, α = .05/6 =.008). The applied α level for each statistical analysis is provided with the results Tables. To determine whether the *t*-test were impacted by between-group differences in sleep, depression and anxiety, we conducted additional *t*-tests analyses using residual values after regressing out global scores on the PSQI, BAI and BDI.

To explore the relationship between subjective and objective measures of memory between groups, we conducted two *post hoc* analyses. First, we explored the hypothesis that mothers’ subjective ratings of their memory may be inaccurate by comparison to their objective scores. We refer to this as ‘memory fidelity’. In our second *post hoc* analysis, we explored the hypothesis that mothers with lower psychosocial wellbeing judge their memory performance more negatively than mothers with better wellbeing. To investigate these *post hoc* hypotheses, we created summary scores of objective memory and wellbeing. For the *objective memory* summary score, we conducted a principal components analysis (PCA) of the measures of objective memory performance (HVLT recall, Digit Span Forward, and Digit Span Backward). For the *wellbeing* summary score, we conducted a separate PCA on the three psychosocial measures (PSQI, BAI and BDI). These two PCAs were conducted in Matlab 2019b, using a singular value decomposition algorithm (‘pca’ function). Based on Kaiser’s recommendation, PCs with eigenvalues >1 were retained for further analysis^44^. Retained PCs were then correlated with global scores on the Prospective and Retrospective Memory Questionnaire (PRMQ). The alpha level for the objective memory and wellbeing analyses were corrected for the number of comparisons (mothers and non-mothers) within each analysis; α =.05/2 = .025. Differences in correlations found for mothers and non-mothers were assessed using Fisher’s z^45^.

## Results

Detailed demographic information is given in Table 1. Mothers and non-mothers did not differ in age or education. Mothers showed significantly worse sleep and higher depression than non-mothers. We calculated menstrual cycle phase for a subset of our sample (*N*=45; 24 mothers and 22 nonmothers), who were naturally cycling, and experiencing a regular cycle length, shown in Supplementary Table 1.

**Table 1:** Demographic information for Mothers and Non-Mothers, including independent samples t-test comparisons between groups for age, education, sleep, depression and anxiety.

|  | Mean(SD)<br>Mothers | Mean(SD)<br>Non-Mothers | *t*-value | Cohen's *d* | *p*-value |
| --- | --- | --- | --- | --- | --- |
| *N* | 43 | 43 | - | - | - |
| Age (years) | 32.5 (3.3) | 31.4 (4.0) | 1.43 | 0.31 | .16 |
| Education (years) | 17.9 (2.6) | 17.6 (2.6) | 0.64 | 0.14 | .53 |
| Sleep (Pittsburgh Sleep Quality Index) | 7.4 (3.2) | 4.7 (2.5) | 4.37 | 0.94 | <.001 |
| Depression (Beck Depression Inventory) | 8.4 (5.5) | 4.9 (4.3) | 3.63 | 0.78 | <.001 |
| Anxiety (Beck Anxiety Inventory) | 11.0 (7.8) | 8.3 (6.4) | 1.32 | 0.28 | .19 |
| Hormonal Contraception (None/ Pill/ IUD/ Implant) | 34 / 6 / 2 / 1 | 26 / 11 / 3 / 3 | - | - | - |
| Months Postpartum | 11.6 (1.7) | - | - | - | - |
| Conception (Natural/IVF/Insemination) | 36 / 6 / 1 | - | - | - | - |
| Multiple Pregnancy (Singleton/Twins) | 42 / 1 | - | - | - | - |
| Delivery Method (Vaginal/Caesarean) | 30 / 13 | - | - | - | - |
| Breastfeeding (Currently/Weaned) | 29/14 | - | - | - | - |
| Total Breastfeeding Duration (Months) | 10.0 (3.8) | - | - | - | - |
| Breastfeeding Duration (Weaned) | 6.5 (4.3) | - | - | - | - |
| Sex of Baby (F/M/Both-Twins) | 22 / 20 / 1 | - | - | - | - |

### Objective & Subjective Cognition

Mothers did not significantly differ from non-mothers on any of the cognitive tasks tested (all p>.05; see Table 2). This result did not change when adjusting for sleep, anxiety and depression (all p>.05). Performance on the cognitive tasks did not show a significant relationship with maternal factors (breastfeeding duration, percentage of time spent in primary care, attachment or self-efficacy; all *p*>.05; Supplementary Table 1). Performance on cognitive tasks did not differ between mothers based on the sex of their child (male/female), or delivery method (vaginal/caesarean; Supplementary Table 2).

**Table 2:** Independent samples t-test comparisons between cognitive performance of mothers and non-mothers, before (raw) and after (adjusted) regressing out the effects of sleep, anxiety and depression. Bonferroni-corrected alpha (.05/6) = .008.

|  | Mean(SD)<br>Mothers<br>(*n*=43) | Mean(SD)<br>Non-Mothers<br>(*n*=43) | *t* | Cohen's *d*<br>(*p*-value) | Cohen's *d*<br>(*p*-value)<br>Adjusted |
| --- | --- | --- | --- | --- | --- |
| Hopkins Verbal Learning Test, Free-Recall | 31.3 (3.1) | 30.7 (3.8) | 0.84 | 0.18 (*p*=.41) | 0.28 (*p*=.21) |
| Hopkins Verbal Learning Test, Delayed-Recall | 11.1 (1.1) | 10.7 (1.5) | 1.46 | 0.32 (*p*=.15) | 0.34 (*p*=.12) |
| Digit Span Forward | 7.1 (1.1) | 7.6 (1.2) | -1.93 | -0.42 (*p*=.057) | -0.32 (*p*=.14) |
| Digit Span Backward | 6.4 (0.9) | 6.8 (1.3) | -1.37 | -0.30 (*p*=.17) | -0.17 (*p*=.43) |
| Symbol Digit Modalities Task | 58.3 (7.4) | 61.8 (10.0) | -1.69 | -0.37 (*p*=.095) | -0.17 (*p*=.44) |
| Reading the Mind in the Films | 66.3 (9.8) | 64.9 (11.5) | 0.87 | 0.19 (*p*=.39) | 0.12 (*p*=.58) |

Mothers self-reported significantly worse subjective memory than non-mothers, *t*(84)=2.40, *p*=.020 and Cohen’s *d=* 0.52. This result was no longer significant when adjusting subjective memory score for sleep, anxiety and depression (Cohen’s *d=*0.19, *p*=.38). All subscale effects also lose significance when adjusting for sleep, anxiety and depression (see Supplementary Table 3).

### Exploring the discrepancy between objective and subjective cognition

We undertook a *post hoc* analysis to explore why mothers and non-mothers showed differences in subjective memory, without significant differences in objective memory. We tested two potential explanations for the incongruence of subjective and objective memory measures for the mother group. First, we considered the possibility that the mother group may be inaccurate in their self-reports of memory, whereas the non-mother group may be more accurate. In other words, this analysis of memory *fidelity* tested whether mothers underestimate their own memory performance on subjective measures. In our second *post hoc* analysis, we considered whether a pessimistic self-report was indicative of negative wellbeing. To this end, we explored the relationship between subjective memory and measures of wellbeing (sleep, anxiety and depression) in mothers and non-mothers.

#### Relationships between Objective and Subjective Measures of Memory

To investigate the relationship between objective memory performance and self-reported subjective memory, we created an *objective memory* summary score by conducting a principal components analysis (PCA) on three measures of memory performance: Hopkins Verbal Learning Test, Digit Span Forward, and Digit Span Backward. Table 3 reports the loading coefficients, eigenvalues, and explained variance of the three obtained principal components (PCs). The coefficients of the first component (explaining 62.38% of the variance) positively load onto all three of the objective measures of memory, with higher values indicating better memory performance (Figure 1a). We then correlated the first PC with the subjective memory score (Prospective & Retrospective Memory Questionnaire [PRMQ]) to investigate the relationship between objective memory performance and self-reported subjective memory.

**Figure 1:**
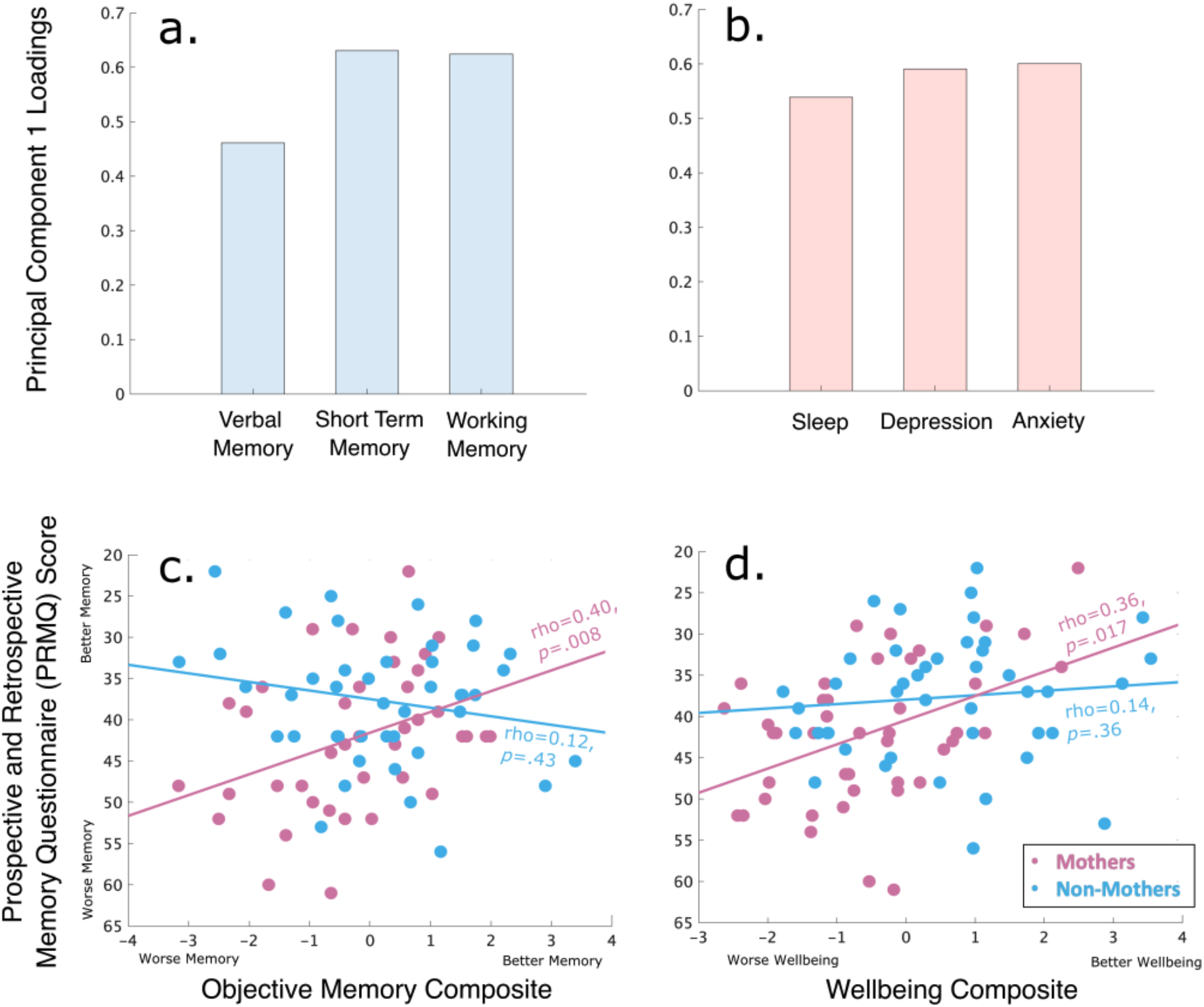
**a)** Component loadings (PCA coefficients) for the first principal component combining verbal memory (HVLT), short-term memory (digit span forward) and working memory (digit span backward) into one objective memory composite score **b)** Component loadings (PCA coefficients) for the first principal component combining sleep (Pittsburgh Sleep Quality Index), depression (Beck Depression Inventory) and anxiety (Beck Anxiety Inventory) into one wellbeing composite score. **c)** Relationships between subjective memory (PRMQ score) and objective memory (first principal component of Hopkins Verbal Learning Task, Digit Span Forward, Digit Span Backward) for mothers (purple) and non-mothers (blue). Higher PRMQ scores indicate worse subjective memory, and higher objective memory values indicate better objective memory performance. For ease of interpretation, we have plotted PRMQ scores from worse (higher score) to better memory (lower score), such that a positive correlation between objective and subjective memory indicates an increase in subjective and objective memory values. **d)** Relationships between subjective memory (PRMQ score) and wellbeing (first principal component of Pittsburgh Sleep Quality Index, Beck Anxiety Inventory and Beck Depression Inventory) for mothers (purple) and non-mothers (blue). Higher scores on the PRMQ depict worse subjective memory, and higher wellbeing scores indicate poorer sleep, and higher levels of depression and anxiety.

**Table 3:**
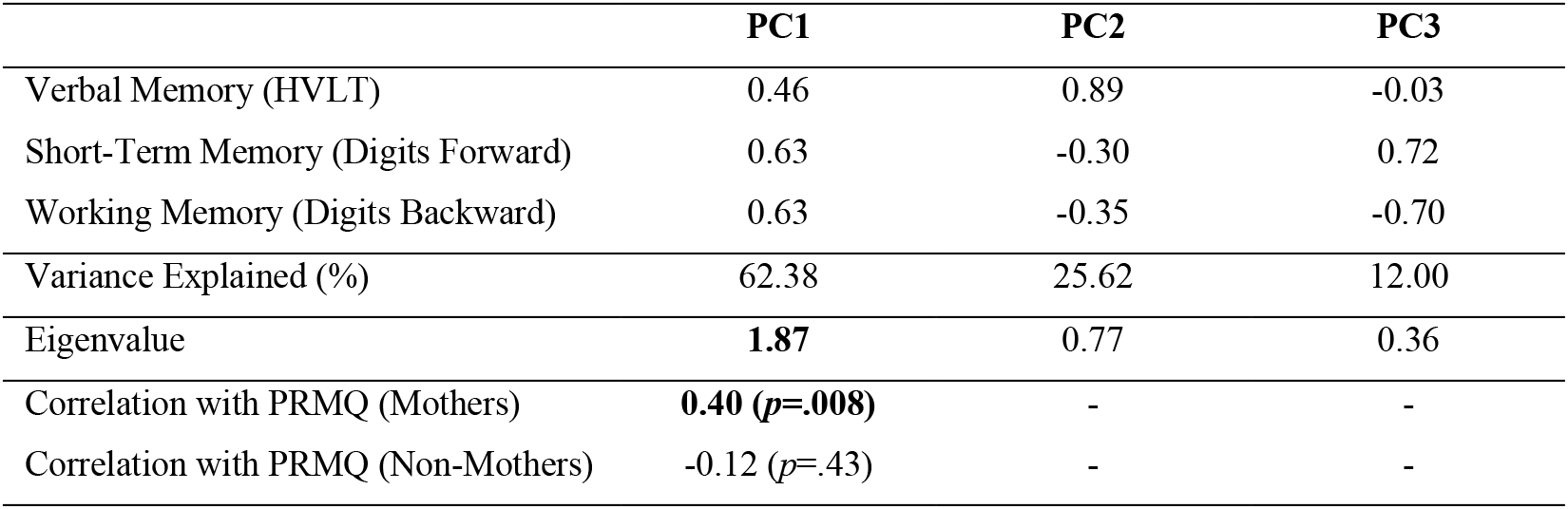
Objective Memory Principal Component Coefficient Loadings, and relationships between Prospective and Retrospective Memory Questionnaire score (PRMQ) and each component for mothers and non-mothers. Alpha = .05/2 = .025

Mothers showed a significant medium positive relationship between subjective and objective measures of memory (*rho*=.40, *p*=.008; Figure 1c, purple), such that mothers who thought (subjectively) that they had better memory, performed (objectively) better. This relationship was consistent across the six subscales of the PRMQ, shown in Supplementary Table 2. No such relationship was obtained between subjective and objective memory for non-mothers (*rho*=-.12, *p*=.43; Figure 1c, blue). These relationships are significantly different from one another (Fisher’s *z*=2.4, *p*=.007), indicating that mothers are more ‘in tune’ with their memory performance than non-mothers.

#### Relationships between Wellbeing and Subjective Memory

To investigate the relationship between wellbeing and subjective memory, we created a summary score of *wellbeing* by conducting a PCA on three psychosocial measures: Pittsburgh Sleep Quality Index, Beck Anxiety Inventory and Beck Depression Inventory. Table 4 reports the loading coefficients, eigenvalues, and explained variance of the three obtained PCs. The coefficients for the first component (explaining 66.73% of the variance) positively load onto all three of the measures of wellbeing (sleep, anxiety and depression), with higher values indicating poorer wellbeing (Figure 1b). We then correlated the first PC with the subjective memory score (PRMQ) to investigate the relationship between wellbeing and subjective memory.

**Table 4:** Wellbeing Principal Component Coefficient Loadings, and relationships between Prospective and Retrospective Memory Questionnaire score (PRMQ) and each component for mothers and non-mothers. Alpha = .05/2 = .025

|  | PC1 | PC2 | PC3 |
| --- | --- | --- | --- |
| Sleep (PSQI) | 0.55 | 0.83 | -0.13 |
| Anxiety (BAI) | 0.60 | -0.28 | 0.75 |
| Depression (BDI) | 0.59 | -0.49 | -0.65 |
| Variance Explained (%) | 66.73 | 19.15 | 14.12 |
| Eigenvalue | **2.00** | 0.58 | 0.42 |
| Correlation with PRMQ (Mothers) | **0.36 ($p=.017$)** | - | - |
| Correlation with PRMQ (Non-Mothers) | 0.14 ($p=.36$) | - | - |

Mothers showed a significant medium relationship between subjective memory scores and wellbeing, such that poorer wellbeing was correlated with worse subjective memory (*rho*=.36, *p*=.017; Figure 1d, purple). Interestingly, the relationship between subjective memory and the wellbeing composite was not significant in non-mothers, (*rho*=.14, *p*=.36; Figure 1d, blue). These relationships were not significantly different from one another (Fisher’s *z*=1.06, *p*=.15).

## Discussion

In this study, we investigated whether there is objective evidence for cognitive decrements in mothers at one-year postpartum. We tested whether mothers one-year postpartum differed from nonmothers on objective and subjective cognitive measures. We hypothesised that mothers would selfreport worse subjective memory than non-mothers, but predicted that mothers would out-perform nonmothers in objective tests of cognition. We obtained significant differences in subjective memory between mothers and non-mothers, in the absence of a difference in objective memory. These differences were no longer significant when we adjusted for differences in sleep, depression and anxiety, suggesting that worse subjective memory is likely impacted by differences in wellbeing. To further explore potential explanations for the incongruence between subjective and objective memory in mothers, we tested two *post hoc* hypotheses; firstly, that mothers show poorer memory fidelity than non-mothers, and underestimate their memory performance on self-report measures. We found a positive relationship between objective and subjective measures of memory in mothers, but not in non-mothers. In the second *post hoc* analysis, we investigated whether mothers with poorer psychosocial wellbeing judge their memory performance more negatively than mothers with better wellbeing. We found a positive correlation between subjective memory and wellbeing, where better wellbeing was associated with better subjective reports of memory. This relationship was not obtained in the non-mother group. Our results challenge the existence of an objective cognitive deficit in mothers at one-year postpartum.

### Objective Cognition

Mothers and non-mothers did not differ on any objective measure of cognition: verbal episodic memory (HVLT), working memory maintenance (digits forward) or manipulation (digit backwards), psychomotor speed (SDMT), or theory of mind (RMFT). These results did not change when adjusting for between-group differences in sleep, anxiety and depression. These results are consistent with previous studies of cognition in the postpartum period, which also report no differences between mothers and controls on measures of short-term memory^6,7,18^, working memory^6,8,11,14,18–20^, and processing speed^5,10,11,14,22^. Notably, the current study provides the first comparison of theory of mind task performance between mothers and non-mothers at any timepoint across pregnancy and the postpartum period, with no differences in performance between groups.

The meta-analysis by Henry and Rendell ^4^ showed subtle verbal memory deficits in mothers in the early postpartum. Extending these results, we found no differences in verbal memory (free- or delayed-recall) between mothers and non-mothers. Our findings are consistent with more recent studies^11,12,14,17^. Glynn ^20^ and de Groot, et al. ^22^ both found verbal memory deficits in primiparous (first-time mothers) and multiparous postpartum mothers (Glynn: three months postpartum; de Groot: eight months postpartum). However, interpretation of memory decrements in these studies was complicated by the fact that both studies also included parous women in their control groups (Glynn: 30% of control group were mothers; de Groot: 58% of control group were mothers), and did not control for group differences in sleep or mood. Thus, on the basis of those results, it remains possible that the obtained cognitive decrements merely reflected poorer cognitive performance by the mothers in the early postpartum, where sleep and mood disturbances are common^46,47^. Indeed, the use of ‘non-pregnant’ control groups, which include women who are mothers, rather than ‘non-parous’ groups, may contribute to the mixed findings in the literature. Studies of cognition in rodents suggest that previous reproductive experience confers cognitive benefits which are long-lasting, perhaps even permanent^2,48^. In humans, reproductive experience is associated with life-long changes to brain structure^49–53^ and function^54^, and improved cognition, including verbal memory^52,53^. Verbal memory also improves longitudinally across the postpartum period (3-12 months), with postpartum mothers showing more evidence of semantic clustering (recalling related words together) than when they were pregnant^15^. Mickes et al.^15^ argued that mothers may increase the use of organisational strategies to support verbal memory, suggesting a potential cognitive mechanism for verbal memory recovery in the postpartum period.

Most studies of postpartum cognition also do not control for poor sleep, depression and anxiety, which are known to be both more prevalent in early parenthood^46,47^ and related to poorer cognitive outcomes outside of a parenthood context^55^. Studies also vary in the time postpartum that mothers are tested, varying from a few days^9^ to two-years after birth^30^. The first weeks and months following birth involve considerable physical, hormonal, and psychosocial changes and rapid adjustment, which stabilise in later postpartum months^28,29^. So, it is likely that the presence and magnitude of cognitive effects of motherhood are influenced by the time of testing postpartum. This argument is compatible with evidence from neuroimaging studies, which suggest that structural brain changes associated with the early postpartum period are variable in direction and magnitude across the transition to motherhood^56–61^. On the basis of extant evidence, it appears that structural reorganisation that occurs during pregnancy and at least the first six-months postpartum, appears to stabilise by two-years postpartum. Here, we show that cognitive differences between mothers and non-mothers are not significant by one-year postpartum, and that sleep and mood are not playing a driving role in objective cognition by one-year postpartum.

Our results are consistent with longitudinal studies of postpartum cognition. Christensen, et al. ^11^ conducted the largest, and most rigorously controlled study of postpartum cognition to date; a prospective longitudinal study, following women before conception, to pregnancy, and into the postpartum period. Of the 806 women tested at baseline, 76 women were pregnant at one of the two follow-up sessions (four- and eight-years later), 188 had become mothers, and 542 had remained nulliparous. No significant differences on any cognitive task were found as a function of pregnancy or motherhood. We conclude that the mixed results in the postpartum cognitive literature may be due to three relevant methodological factors; 1) some studies include parous women in their control groups, 2) some studies control for the effects of sleep, depression, anxiety, 3) studies vary in the time postpartum that mothers are tested, 4) laboratory tasks may have less ecological validity than naturalistic tasks, and 5) studies may find different results if they tested parenting specific skills and abilities.

### Subjective Memory

Despite showing no significant differences from non-mothers on objective measures of memory or cognition, mothers self-reported significantly worse memory than non-mothers. These findings support our hypotheses, and are consistent with previous studies^7,8,14,20,31^ and meta-analyses^4^ showing subjective memory deficits in the postpartum period. In the present study, and others^7,8,12,14,31,32^, significant differences between mothers and non-mothers in subjective memory are present in the absence of any objective memory differences. This discrepancy suggests that mothers are consistently reporting perceived memory deficits without an objective memory deficit. This discrepancy begs the question - *why* do mothers and non-mothers show differences in subjective, but not objective memory? If the difference in subjective memory isn’t because mothers are recalling less, then what is driving this consistent self-perceived deficit? One possible interpretation is the *fidelity* of self-reported memory – whether subjective reports of memory performance align with objective performance.

Mothers showed a significant positive relationship between subjective and objective measures of memory, such that better self-reported memory was associated with better objective memory performance. No such relationship was found between subjective and objective measures of memory in non-mothers. In other words, mothers’ subjective ratings of their memory were aligned with their objectively measured memory scores, whereas non-mothers showed poor alignment between subjective and objective memory. Here, we use the term ‘memory fidelity’ to refer to the alignment between subjective and objective memory. While it may initially seem counter-intuitive that mothers report worse memory, but are more ‘in tune’ with their memory performance than non-mothers, close scrutiny of Figure 1c confirms that regardless of whether objective memory performance was high or low, nonmothers subjectively rated their memory as high, whereas this was not the case for mothers. Similar positive relationships between subjective and objective measures of memory have been reported in pregnancy^62–64^, which have been interpreted as pregnant women accurately reporting memory deficits. Subjective memory fidelity in mothers may be a consequence of increased self-awareness in motherhood^3,65^.

The second explanation that we explored to account for mothers’ worse subjective memory is poorer maternal wellbeing. We hypothesised that mothers who experienced poorer wellbeing (poorer sleep, higher symptoms of depression or anxiety) might be more inclined to think negatively of their memory performance, and report worse subjective memory. We found a relationship between mothers’ self-reported memory and wellbeing, where poorer wellbeing was associated with worse subjective memory ratings. Similar relationships between subjective memory and disturbances in sleep and mood have been shown in the first year postpartum^6–8,14^, and in the transition to menopause^67^.

We speculate that mothers’ increased awareness of their memory performance may arise from three potentially interacting factors; a greater daily cognitive load (i.e., an increase in the number of items and tasks to remember in daily life), an increased attention to memory lapses, and poorer wellbeing. The postpartum period is a time of rapid behavioural adaptation, with novel challenges, tasks and patterns to attend to and recall. For example, leaving the house with a one-year-old in tow, requires a new mother to remember a host of items – nappy bag, dummy, a favourite toy, snacks, a change of clothes, a bottle – the list goes on. With an increased cognitive load, there become more opportunities to forget an item or task. Memory errors in the postpartum period also carry heavier consequences (e.g., an unsettled baby, a sleepless night, in the extreme – harm to the baby), making memory lapses more salient. Pregnancy and early motherhood may make women more attuned to their functioning, through a higher sensitivity to minor memory or concentration lapses, which perhaps would have otherwise been ignored, or considered inconsequential before pregnancy^65^. We speculate that the increased cognitive load of daily life in early motherhood represents a cognitive challenge, with higher consequences for memory lapses, which encourages mothers to evaluate and re-evaluate their subjective memory regularly, making them more in tune with their memory performance. In contrast, a possible reason that non-mothers do not show a similar relationship between their subjective memory and their objective memory performance is that they are not experiencing, or not aware of, memory lapses in their daily lives. We may see higher memory fidelity in mothers because they are being cognitively challenged, and they are more aware of their memory errors when they occur. This continuous reevaluation may also be related to the societal expectation of a cognitive decline in motherhood^66^, contributing to a confirmation bias towards expected cognitive changes in pregnancy and the postpartum period. The mismatch between subjective and objective memory is also related to mood disorders outside of the postpartum period, where people with more severe depression, anxiety, and general psychological distress bipolar disorder rate their memories as more impaired^68–70^, even after controlling for differences in objective cognitive performance^71^. Additionally, people with mood disorders also show increased memory fidelity, with a greater sensitivity to differences in their memory ability, and more ‘insight’ into cognitive changes^68–70^. Taken together we speculate that maternal memory perception is influenced by increased cognitive load, a greater awareness of memory lapses, and poorer wellbeing.

Our results further suggest that measuring and controlling for differences in sleep, depression, and anxiety is essential when assessing cognition in the postpartum period, and providing appropriate support for new mothers who are experiencing poor wellbeing. To determine whether the mechanism behind subjective memory decline in mothers is specific to parenthood, or sleep, depression and anxiety, future studies should compare mothers to a nulliparous control group matched for these factors.

### Future Directions

Whilst we did not observe any differences in objective cognitive performance between mothers and non-mothers, we cannot rule out the possibility that motherhood may induce cognitive changes, apparent to mothers themselves, but so subtle that they do not affect current laboratory-based tests. It is also possible that mothers experience subtle cognitive deficits in their everyday lives, which are not evident in the idiosyncratic environment of cognitive test conditions. For example, previous studies in pregnancy have found memory deficits in tasks completed in the mother’s home environment, when performance on laboratory tasks was intact^16,64^. Furthermore, we cannot rule out the possibility that mothers may show cognitive differences to non-mothers in other tests of objective memory. Tasks with greater ecological validity, measured in a mother’s home, and perhaps including baby-related information, may uncover differences in cognition that we did not measure here.

Furthermore, it is important to note the cross-sectional design of this study when interpreting the results. The cross-sectional design means that we can only interpret *differences* between mothers and non-mothers, not *changes* across pregnancy and the postpartum period^72,73^. This also means that we cannot rule out the possibility that mothers in our sample experienced an objective decline compared to their own pre-pregnancy baseline performance. However, our results are consistent with well-controlled longitudinal studies, which also found no differences in objective cognition between mothers and nonmothers across pregnancy and the postpartum period^11,14,17^.

Cognitive changes in early motherhood are likely related to the extreme hormonal fluctuations of pregnancy, parturition, and lactation. However, it is also possible that mothers at one-year postpartum are not experiencing measurable memory deficits, but instead self-report forgetfulness which they perceive due to cultural expectation and anecdotal experience of the ‘baby-brain’ stereotype. In Australia, there is a strong societal narrative regarding baby-brain and its impact on maternal cognition^1,11,74^. Crawley, et al. ^75^ argue that the pregnancy-related decline in memory performance has been exaggerated in the scientific literature, and that subjective reports of memory difficulties result from cultural expectations of a cognitive deficit in pregnancy. These expectations are perpetuated by media, pop culture, and even in the advice of medical caregivers^76^. The baby-brain stereotype may be so established and internalised that it becomes a somewhat ‘self-fulfilling prophecy’, where mothers’ perception of their memory are seen through the lens of expected deficits^12,32^. For example, an everyday lapse in memory may be falsely attributed to postpartum status because of the context provided, and is then perpetuated by confirmation bias^76^, where evidence in support of forgetfulness is disproportionately attended to, remembered, and discussed. This cycle works to incorrectly verify ‘baby-brain’, maintaining the stereotype that motherhood is accompanied by memory decline. It is crucial that new and expectant mothers receive accurate information to appropriately guide their self-perception in the transition to motherhood.

## Conclusion

We show significant differences in subjective memory between mothers and non-mothers, in the absence of objective differences. In interpreting the discrepancy between subjective and objective memory in the postpartum period, we found evidence that mothers have greater self-reported memory ‘fidelity’, suggesting mothers may be more in tune with their memory performance than non-mothers, and that mothers with poorer wellbeing (sleep, anxiety, and depression) perceive worse subjective memory. Our results, alongside past literature, challenge the existence of cognitive deficits at one-year postpartum. Our results have implications for how new mothers are perceived by society, and how they think about themselves and their own cognitive ability. Especially when subjective perception of cognitive ability is so strongly affected by expectation, it is critical that we do not overestimate, exaggerate, or extrapolate deficits in pregnancy to the postpartum period. Regardless of whether objective memory deficits persist into the postpartum, discussion of memory issues may serve as an opportunity to seek and receive additional support from health-care professionals. Memory complaints in the postpartum must therefore not be dismissed, especially as they may indicate difficulties with poor sleep, or low mood. New and expectant mothers must receive accurate information to appropriately guide their self-perception in the transition to motherhood. In future, health-care providers should explain these findings to new and expectant mothers, that cognitive deficits in the postpartum period are not inevitable, and instead focus on providing appropriate care and support for mothers who express difficulties with poor sleep, or low mood.

## Acknowledgements

The authors are supported by the Australian Research Council (ARC) Centre of Excellence for Integrative Brain Function (CE140100007), and this study received support from an early career researcher project grant from the Centre to ERO. SDJ is supported by an Australian National Health and Medical Research Council Fellowship (APP1174164).

## Availability of data

All data supporting the results of this study are presented in the tables and supplements of the manuscript.

## Supplementary Information

**Supplementary Table 1:** Menstrual Phase for a subset of the sample who were naturally cycling and experiencing a regular cycle length with menstrual periods.

|  | Mothers (N) | Non-Mothers (N) |
| --- | --- | --- |
| Early Follicular | 5 | 1 |
| Late Follicular | 4 | 6 |
| Early Luteal | 5 | 8 |
| Late Luteal | 10 | 7 |

**Supplementary Figure 1:**
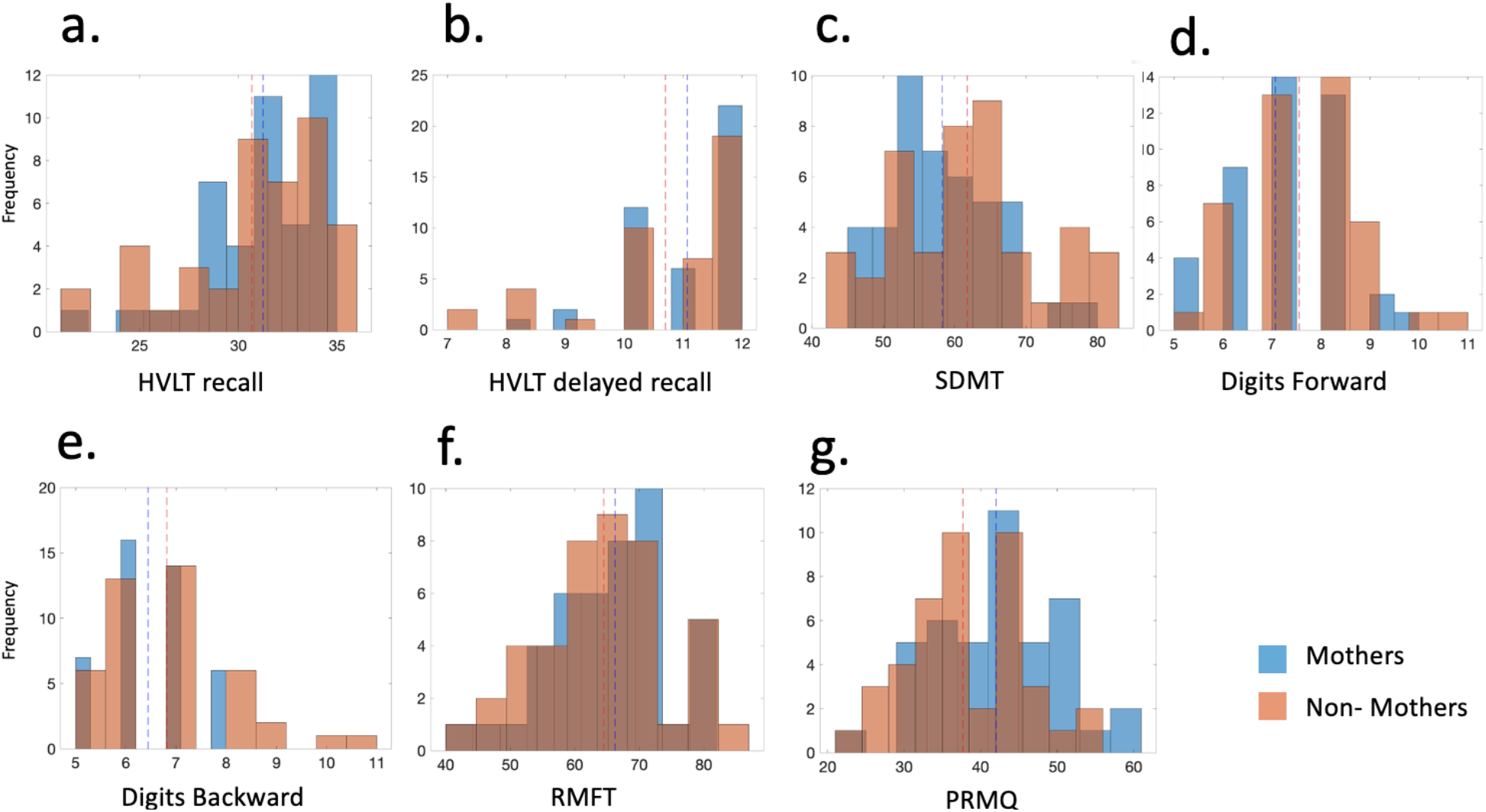
Histograms for the raw scores for the six cognitive tasks, and subjective memory questionnaire. Mothers (in blue), non-mothers (in red), vertical dashed lines show the mean for each group.

**Supplementary Table 2:** Correlations between Subscales of the Prospective and Retrospective Memory Questionnaire, with the Objective memory composite score for mothers and non-mothers

|  | Mothers |  | Non-Mothers |  |
| --- | --- | --- | --- | --- |
|  | Spearman's Rho | p-value | Spearman's Rho | p-value |
| Total PRMQ (reported in Manuscript) | .40 | .008 | -.12 | .43 |
| Prospective | .35 | .02 | -.11 | .50 |
| Retrospective | .38 | .012 | -.09 | .55 |
| Long term | .43 | .005 | -.13 | .39 |
| Short term | .33 | .03 | -.08 | .60 |
| Self-cued | .43 | .004 | -.18 | .26 |
| Environmental-cued | .30 | .05 | -.02 | .91 |

**Supplementary Table 3:**
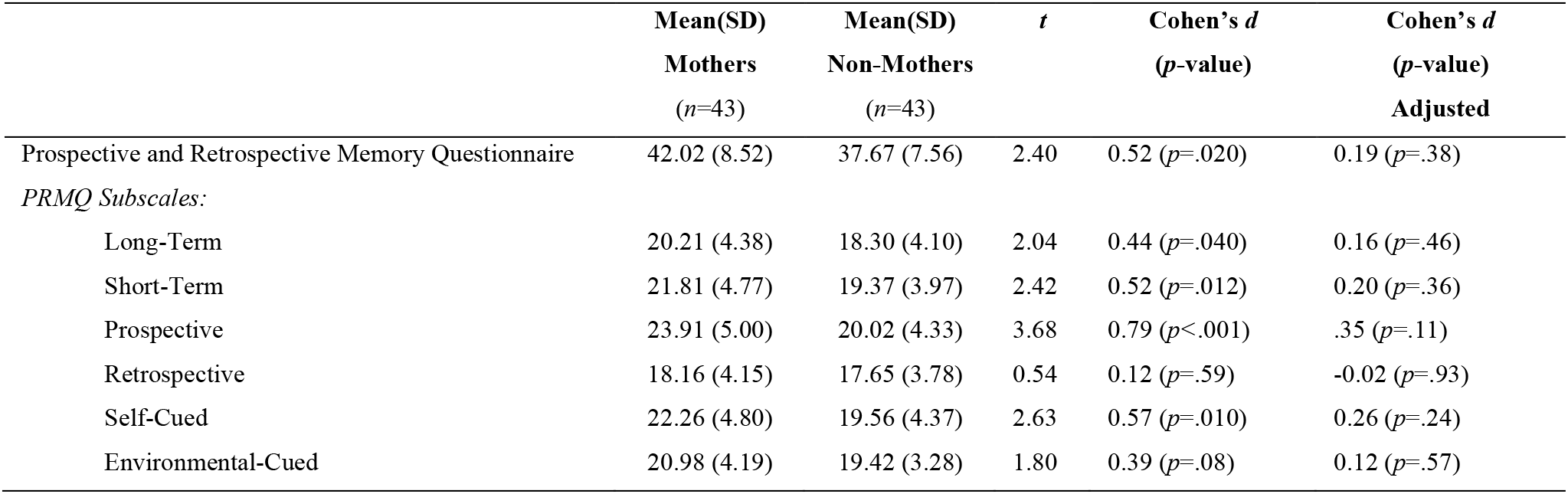
Independent samples t-test comparisons between subjective memory ratings of mothers and non-mothers, before (raw) and after (adjusted) regressing out the effects of sleep, anxiety and depression.

